# Early life stress of maternal deprivation and peer-rearing jeopardize mesoprefrontal and mesolimbic dopamine receptors in the rhesus monkey

**DOI:** 10.64898/2026.02.28.708755

**Authors:** Sally B. Seraphin, M. Mar Sanchez

## Abstract

Early life stress (ELS) in primates alters dopamine function, contributing to addiction, hyperactivity, cognitive deficits, aggression, and social subordinance. To assess whether dopamine receptor densities are affected by ELS, male juvenile rhesus monkeys (Macaca mulatta) were either mother-reared (MR, N=6) in a semi-natural environment or nursery-reared (NR, N=6) with peers in a laboratory. At 1 ½ years of age, subjects were sacrificed and the left prefrontal cortex (PFC), striatum (caudate and putamen), nucleus accumbens (NAcc), and claustrum (CLA) were explored through quantitative autoradiographic studies of dopamine receptor-1 (DRD1) and -2 (DRD2) conducted using [125I]-(+)-SCH 23982 and ^125^I-Epidepride, which have high affinity and selectivity for DRD1 and DRD2, respectively. No group differences emerged in striatal or NAcc receptor binding. However, MR monkeys exhibited significantly greater DRD1 binding in the left orbital PFC and significantly greater DRD2 binding in both the left medial PFC and right CLA compared to NR. These findings implicate the medial PFC (stress vulnerability, cognition), orbital PFC (reward valuation), and CLA (anxiety modulation) as critical sites disrupted by maternal deprivation. Therefore, we propose that nursery-rearing induces a hypodopaminergic prefrontal-claustral ecophenotype, underlying the cognitive, affective, and social impairments observed in NR monkeys.

## INTRODUCTION

### Attachment, Social Bonding and Dopamine

For over a century, attachment and bonding have preoccupied animal researchers. Their resulting scholarship spans a broad continuum from the classical ethology studies of Konrad Lorenz, Niko Tinbergen, and Karl von Frisch (Font, 2023; Hinde, 2005), to the early psychological studies of John Bowlby, Mary Ainsworth, and Harry Harlow (Vicedo, 2010), to the recent cellular molecular neurobiological approaches established by Larry Young (Liu and Murphy, 2024). Consequently, we know social affiliation is essential for species living in communities that require cooperation for survival and reproduction. Bonding is especially vital for altricial animals, where the prolonged dependence on parental care is positively associated with neuroplasticity (Gómez-Robles et al., 2024; Lauby et al., 2021). Dopamine (DA) directs the formation of bonds and attachments by incentivizing reward (Blumenthal and Young, 2023; Zeevi et al., 2021). That social and affiliative behavior are coordinated by DA and oxytocin (OT) is evidenced by the presence of DA receptors on oxytocinergic neurons within the hypothalamus (Baskerville and Douglas, 2010). DA participates in the integration and interpretation of environmental information that is relevant for executing the behavioral adjustments necessary for survival, reproduction, and evolutionary adaptation.

DA so fundamentally orients the individual towards important objects of their attention and sources of reward, that the connection between a mated pair, or its evolutionary antecedent the mother-infant bond, can be characterized as addictive (Burkett and Young, 2012). Indeed, the progression from substance abuse to addiction is mediated by OT, DA, and glucocorticoid systems (Kim et al., 2017). In a study of partner preference among monogamous prairie voles (*Microtus ochrogaster*), mating caused 50% increased extracellular DA in the nucleus accumbens (NAcc) (Gingrich et al., 2000). Moreover, using microinjection, the resulting partner preference was disrupted by the Dopamine Receptor-2 (DRD2) antagonist eticlopride and restored by the DRD2 agonist quinpirole (Gingrich et al., 2000). Conversely, the experience of yearning after reminders of the deceased beloved is associated with increased activity in craving and reward-related areas, such as the NAcc, in people suffering from complex grief (O’Connor et al., 2008). Finally, repeated maternal separation has been shown to increase the DA response to stress, lower levels of dopamine transporter (DAT, the rate limiting enzyme in DA synthesis), and alter the locomotor behavioral sensitivity to both stress and cocaine or amphetamine (Meaney et al., 2002).

Reliable, nurturing, and attentive care is key to the success of social animals. For mammals in particular, early environmental experiences determine the quality of attachment, which serves as a foundation for self-regulation and social behaviors. Behavioral anomalies associated with early adverse experiences call to mind the features of psychological disorders and neurological conditions involving DA neurotransmission. Early social deprivation, abuse, or neglect have been linked to social impairment, affective dysregulation, elevated addiction risk, and cognitive deficits (Dettmer and Chusyd, 2023; Fahlke et al., 2000; Parise et al., 2025; Sanchez et al., 2001). For instance, in keeping with their altered amygdala function, non-human primates exposed to early stress, such as social-isolation (Harlow et al., 1965), repeated separation from a caregiver (Harlow and Suomi, 1974), or parental abuse/neglect (Fahlke et al., 2000; Maestripieri et al., 2000; Maestripieri, 1994; Suomi, 2005, 1997) are more fearful and aggressive. Increased self-injurious behavior, involuntary or stereotypic movements; impaired attention and memory; and abnormal social behavior or communication have also been observed in animal models of maltreatment (Dettmer and Chusyd, 2023; Sanchez et al., 2001; Seraphin et al., 2022). In short, the eco-phenotype typically associated with early life stress (ELS) may partly emerge from developmental programing for an environment that is devoid of stable attachment, plagued by scarcity, and generally marked by severe uncertainty.

### Dopamine Receptor Function & Neuroanatomy

In keeping with its global significance for behavior, dopaminergic signaling is ubiquitous in the central and peripheral nervous systems (Luo and Huang, 2016). Brain DA systems have emerged as a factor in the etiology of many neurological maladies, including disorders of cognition, affect, aging, addiction, and even reproduction.

Nevertheless, the role of DA is neither nonspecific nor homogenous. Instead, its discrete functions are rooted in cell bodies primarily localized in the midbrain (i.e., ventral tegmental area (VTA), substantia nigra pars compacta (SNpc), retrorubral field (RRF)), and to a lesser degree, hypothalamic and thalamic regions (i.e., posterior hypothalamus, arcuate nucleus, and zona incerta)(Bolis et al., 2001; Klein et al., 2019). Further, DA’s broad influence unfolds through extensive axonal projections spanning four neuronal pathways, where its levels are modulated through uptake or cellular absorption by DAT, and its actions are mediated through activation of five different DA receptors (Beaulieu and Gainetdinov, 2011; Luo and Huang, 2016). The presence of these receptors –in such far flung brain regions as the cerebellum, expand DA’s reach and behavioral significance (Flace et al., 2021).

*Dopamine Receptors* - DA receptors belong to the G-protein-coupled receptor family and are classified as DRD1-like (DRD1, DRD5) or DRD2-like (DRD2, DRD3, DRD4) (Beaulieu and Gainetdinov, 2011; Neve et al., 1997). They can be functionally differentiated by their effect on adenylate cyclase activity, with the DRD1-like receptors *stimulating* and DRD2-like receptors *inhibiting* its action. Thus, they may be viewed as having opposing roles in intracellular signaling with DRD1-like receptors *increasing* and DRD2-like receptors *decreasing* the production of cyclic adenosine monophosphate (Cyclic AMP, cAMP) (Bolis et al., 2001; Di Chiara et al., 2014; Kim et al., 2022; Ophuis et al., 2014; Wooten, 2001). To support their distinct behavioral functions, the neuroanatomical distribution of DRD1-like and DRD2-like receptors is region-specific (Choi et al., 1995; Fujimoto et al., 2025; Meador-Woodruff et al., 1991; Sung Choi et al., 1995). These behavioral effects are defined by four main, functionally segregated DA pathways (Bergson et al., 1995; Terauchi et al., 2025) controlling movement, cognition, reward, and basic drives.

*Brain Dopamine Pathways* - DA neurons project independently from the ventral tegmental area (VTA) to distinct brain regions, forming “projection-defined” circuits with specialized roles (Vander Weele et al., 2019). The nigrostriatal pathway, originating in the substantia nigra (SN) and projecting to the striatum, regulates motor control and is implicated in Parkinson’s and Huntington’s diseases (Mishra et al., 2025). The mesocortical pathway links the VTA to the frontal cortex, supporting cognition, memory, and attention. Its dysregulation contributes to ADHD, Alzheimer’s disease, PTSD, depression, and negative symptoms of schizophrenia (Lyons et al., 2000; Mathew et al., 2001; Neve et al., 1997; Nieoullon, 2002). The mesolimbic pathway, connecting the VTA to the nucleus accumbens (NAcc) and amygdala, governs emotion, reward, fear, and addiction (Brandão and Coimbra, 2019; Jin et al., 2023; Mehr et al., 2025). Unlike the mesocortical circuit, which responds strongly to aversive stimuli, the mesolimbic system is reward sensitive. Cortical fibers linking the frontal cortex with subcortical claustrum (CLA) and amygdala also regulate anxiety and stress behaviors (Niu et al., 2022; Wang et al., 2023). Overlapping mesocortico-limbic circuits are implicated in depression and autism spectrum disorder (ASD), where DA dysfunction underlies social and repetitive deficits (DiCarlo and Wallace, 2022). Finally, the tuberoinfundibular pathway (TIDA) connects the hypothalamus to the pituitary, controlling prolactin, oxytocin, and steroid release (Giuliano and Allard, 2001; Melis et al., 2022; Qi-Lytle et al., 2023). As such, it is involved in both antisocial and prosocial behaviors ranging from aggression and violence (Rosell and Siever, 2015) to trust and maternal care (Georgescu et al., 2022; Koyama et al., 2024). For example, in a study of reproductive neurocrines in cultured hypothalamic explants from the marmoset monkey (*Callithrix jacchus*), DA secretion was lower while OT and PRL secretions were higher in parentally experienced males compared to naive males (Woller et al., 2012).

### Early Life Stress and Dopamine

The epigenetic control of dopaminergic neuroadaptations to ELS has been explored through experiments utilizing prenatal and postnatal paradigms in an array of species. The nature of clinical outcomes depends on the timing of early life adversity and connected critical or sensitive periods of development (Parise et al., 2025). For instance, prenatal stress in rhesus macaque infants is associated with increased cerebrospinal fluid (CSF) levels of two catecholamine metabolites (DOPAC: 3,4-dihydroxyphenylacetic acid and MHPG: 3-methoxy-4-hydroxyphenylglycol), at 8 and 18 months of age (Schneider et al., 1998), and increased striatal DRD2 binding potential at 5 to 7 years of age (Roberts et al., 2004). As a model of early and chronic psychosocial stress, social isolation (SI) rearing often results in the most debilitating outcomes and yet consistently results in altered DA functioning. For instance, both structural and functional mesocorticolimbic DA system modifications (e.g., dendritic reduction) have been documented in behaviorally impaired SI rats (Wang et al., 2012), which have a tendency for increased anxiety like behavior and drug or alcohol self-administration (Yorgason et al., 2013). Across numerous rat studies, SI has been linked to increased functional high affinity state DRD2 in the striatum (Fone and Porkess, 2008; King et al., 2009), elevated DOPAC in the shell of the NAcc (Bortolato et al., 2011), increased DAT activity in the NAcc core (Yorgason et al., 2013), and increased DA release in both the in the shell (Bortolato et al., 2011) and core of the NAcc (Yorgason et al., 2013). That elevated DA or DOPAC were not observed in the medial prefrontal cortex (mPFC) (Bortolato et al., 2011) of SI rats suggest that compensatory adjustments to DA signaling are region specific.

SI has also been extensively modeled in non-human primates. In a study of adult male and female rhesus monkeys (Macaca mulatta), animals previously deprived of social contact and reared in isolation for the first 9 months of life had lower caudate and putamen levels of tyrosine hydroxylase (TH, the rate limiting enzyme in DA synthesis) in axonal fibers compared to controls (Martin et al., 1991). Moreover, the loss of TH immunopositive fibers was paralleled by a general reduction of TH immunopositive neurons in the SN (Martin et al., 1991). Although a quantitative analysis of TH within the TIDA of adult rhesus monkeys, observed no differences between SI animals and controls leading authors to speculate that the lacking effect of rearing can be attributed to neuronal cell loss due to aging (Ginsberg et al., 1993), this may be another instance of region specificity.

Institutionalization is a classic model for dysfunctional caregiving and its lasting effects on attachment. In a comparison of 6-36 month old children in an orphanage with peers residing with family, the orphans were found to have greatly increased anxiety, decreased blood DA and serotonin (5HT), and increased blood norepinephrine (NE) compared to their family-housed counterparts (Gogberashvili et al., 2011). Further implicating DA, motivational deficiencies are common outcomes of institutionalized rearing. For instance, in a study using monetary rewards to compare former Romanian orphans once exposed to institutional deprivation with non-adopted British counterparts, low anticipatory responses were linked to diminished recruitment of the striatum in the former orphans (Mehta et al., 2010). Peer-rearing (PR) resembles institutionalization because both are models of social-integration without consistent parental care. In addition to hypothalamic–pituitary–adrenal (HPA) axis dysregulation, non-human primate studies of peer-rearing also demonstrate altered CSF levels of OT, DA, NE, 5HT, and their metabolites compared to controls . In contrast to mother-reared (MR) rhesus monkeys, we have observed lower 5HT, reduced dopaminergic tone, and lacking correlations between CSF monoamines, catecholamines, and their metabolites among peer housed, nursery-reared (NR) monkeys (Seraphin et al., 2022).

Rodent and non-human primate studies show that neurobiological effects of ELS can manifest in both structural and functional changes in DA circuits. Interestingly, these stress-related modifications include both inflated and diminished dopaminergic signaling, in keeping with both nature and timing of insult. For example, maternal separation is linked to increased DA turnover in guinea pigs (Tamborski et al., 1990), as well as both decreased DAT and elevated DA in rats (Meaney et al., 2002). Nevertheless, maternal deprivation and separation paradigms consistently reveal changes to neurotransmitter functions, including dysregulation of DA neurons. For example, whole-cell patch clamp recording on rat pup midbrain sections revealed that merely one 24-hr period of maternal deprivation on postnatal day 9 is sufficient to epigenetically weaken both excitatory and inhibitory synapses, or long-term depression (LTD), in DA neurons of the VTA (Authement et al., 2015). By decreasing the GABAergic inhibition of DA neurons, this potentially diminishes cell firing and release of DA (Dacher and Nugent, 2011; Hanson et al., 2021). These rodent experimental outcomes are echoed in one human study where adults with childhood histories of poor parental care showed elevated mesoaccumbal DA, and enhanced behavioral responsivity to reward after treatment with the DA agonist methylphenidate (MPH) (Engert et al., 2009). When similarly challenged by repeated separations, elevated levels of Homovanillic Acid (HVA, a DA metabolite) and reduced levels of MHPG, have been observed in peer-reared rhesus monkeys, compared to MR controls (Wood et al., 2021). Using an exceptional ecologically valid approach, Coplan and colleagues discovered altered DA functioning as an outcome of acute environmental uncertainty. In studies of bonnet macaque (*Macaca radiata*) mothers exposed to uncertain, variable foraging demand (VFD) conditions during the early postnatal period, they observed that decreased infant cortisol and HVA resulted from unpredictable, mother-infant interactions caused by foraging stress (Coplan et al., 1998; Meyer and Hamel, 2014). Notably, exposure to deficient maternal care can have lasting consequences such as persistently impaired DA signaling and behavioral responses in the adult. For instance, a quantitative receptor autoradiography study of maternally separated rats compared to non-handled controls found reduced NAcc core and striatal DAT as well as reduced DRD3 binding and mRNA levels in the NAcc shell (Brake et al., 2004). In further comparisons with controls, the maternally separated animals exhibited increased locomotor sensitization to amphetamine when repeatedly stressed, and increased cocaine dose-dependent sensitization of their locomotor response to tail-pinch (Brake et al., 2004).

### Dynamics of Dopaminergic Programing

Although early childhood adversity is a risk factor for developing psychopathological conditions associated with DA dysfunction, such as schizophrenia and addiction, there remains little consensus on the DA mechanisms contributing to sequelae of ELS (Bonapersona et al., 2018). Our understanding of the developmental programing of dopaminergic functions is key to the prevention and mitigation of their adverse effects. Several mechanisms potentially link ELS with DA function:

**1. Amplified DA signaling** - Mesoprefrontal DA neurons are selectively activated by stress. This is supported by the observation of elevated DOPAC levels in the VTA of animals stressed for 10 minutes, and elevated levels of both DOPAC and HVA in the mPFC of animals stressed for 30 minutes by the mere exposure to the responses of foot-shocked rats (Kaneyuki et al., 1991). Stress further alters the functional morphology of the VTA by: a) increasing DA neuron excitability and baseline DA levels in the NAcc (Hanson et al., 2021) and b) enhancing reward-related connectivity between the VTA and mPFC (Hanson et al., 2018) in depressed adults with prior histories of childhood maltreatment. Egerton et al. (2016), observed an association between histories of childhood adversity in young adults and elevated dopamine synthesis capacity that conveys increased striatal dopamine function.
**2. Altered DA turnover -** Acute and chronic stress may differ in their effects on DA secretion and metabolism. The ratio of DOPAC to DA, which is a measure of DA turnover, is attenuated by repeated maternal separation in rat pups (Mathew et al., 2001). TH, as the rate limiting enzyme in DA synthesis, can increase or decrease availability of DA when it is up-regulated or down-regulated respectively. In an in vivo microdialysis experiment, Finlay and colleagues (1995) detected increased extracellular DA in the mPFC under both acute (30 minutes of tail pressure) and chronic stress (3-4 weeks of 5°C cold exposure). However, only acutely stressed animals responded with decreased extracellular DA when treated with the anxiolytic, diazepam (Finlay et al., 1995).
**3. Suppressed DA signaling** – Stress precipitates functional compensatory responses, such as receptor downregulation or desensitization, to support allostasis. Thus, chronic stress may “blunt” or lessen the capacity to synthesize striatal DA, resulting in attenuated responses to rewards and stressors in limbic regions associated with reward processing (Bloomfield et al., 2019). Consequently, Deficits in reward learning have been observed in human adolescents with significant histories of childhood adversity (Hanson et al., 2017). Also, the downregulation of NAcc DRD1, DRD2, and DAT gene expression have been observed in maternally deprived rats that subsequently show impaired spatial navigation ability on the Morris water-maze test (Zhu et al., 2010).
**4. Diminished top-down cognitive suppression of limbic responses** – Stress precipitates *structurally* compensatory responses, such as altered dopaminergic neuronal connectivity, in service of allostasis. For example, specific modifications to adolescent DRD1 and DRD2 PFC microcircuitry were observed in differently reared rats where 4-hrs of daily maternal separation stress, between post-natal day 2-20, culminated in significantly less DRD1 and DRD2 expression on glutamatergic projection neurons linking the PFC and NAcc (Brenhouse et al., 2013). In a study of adolescent female rats, maternal separation increased DRD2 receptor mRNA expression was observed in the prelimbic cortex while elevated TH immunoreactive fibers were found in both the NAcc and prelimbic cortex (Majcher-Maślanka et al., 2017), suggesting a disruptive effect of ELS on DADR’s modulation of cognition and emotion.

Considering what is known about their roles in mediating complex mesocortical-mesoprefrontal and mesolimbic DA driven behavioral processes, we endeavored to explore whether DRD1 and DRD2 levels differ as a function of rearing in rhesus monkeys. In a previous study comparing MR and NR rhesus monkeys, we reported evidence to support suppressed DA signaling in NR. Specifically, through a standard measure of dopaminergic tone, we showed that NR produced insufficient levels of prolactin in response to pharmacological challenge by Raclopride, which is a DRD2 antagonist (Seraphin et al., 2022). When observed in a social setting, indexes of dopaminergic tone in NR monkeys were negatively correlated with increased aggressive, self-injurious, and repetitive behaviors (Seraphin et al., 2022). To corroborate and extend these findings, we hypothesized that the ELS of maternal deprivation and peer-rearing would result in altered densities of DRD1 and DRD2 in the basal ganglia and reduced DRD1 and DRD2 in the PFC of NR compared with NR. In this report, our two-part strategy was as follows:

#### 1. Evaluate suppression of dopamine signaling in NR

While acute stress is associated with increased cortical dopamine release, inescapable stress that is chronic can also down regulate dopaminergic function (Howes et al., 2017). Reduced TH immunoreactive neurons have been reported in the caudate and putamen of socially deprived rhesus monkeys (Martin et al., 1991), suggesting that abnormal behaviors from deprivation alter basal ganglia DA. Since these regions are innervated by VTA DA fibers, we examined DRD1 and DRD2 densities in the nucleus accumbens (NAcc), caudate, and putamen, predicting altered receptor levels in NR compared to MR monkeys.

#### 2. Assess weakened top-down suppression of limbic responses in NR

Being rich in glucocorticoid receptors (Sanchez et al., 2000), the PFC is highly sensitive to stress (Crane et al., 2003; Figueiredo et al., 2003; Finlay et al., 1995; Jedema et al., 1999; Stevenson and Gratton, 2003). Stress elevates cortical NE and DA (Finlay et al., 1995; Jedema et al., 1999), potentially altering DA receptor expression.

While childhood maltreatment results in medial orbitofrontal grey matter loss (De Brito et al., 2013), rodent studies show increased reward sensitivity and impulsivity (Kirkpatrick et al., 2013) as well as downregulation of DRD2 after maternal separation (Mahmoodkhani et al., 2022). Given that NR monkeys display socioemotional and cognitive deficits (Sánchez et al., 1998; Seraphin et al., 2022) similar to human disorders with DA dysregulation (Heinz, 2002), we predicted reduced DRD1 and DRD2 in the PFC of NR monkeys. Such decreases would reflect suppressed DA signaling and help explain their attentional and motivational impairments.

## MATERIALS AND METHODS

### Rearing of Animal Subjects

Twelve male rhesus monkeys (*Macaca mulatta*), residing at the Yerkes National Regional Primate Research Center’s (YNPRC) Main and Field Stations were employed as subjects of this experiment. All animals were born of middle dominance ranking mothers, over a one-month period. For one-year post-partum randomly chosen monkeys were reared by their birth mothers, in a quasi-natural environment, with periodic access to outdoor enclosures at the YNPRC Field Station, in Lawrenceville, GA. At approximately 12 months of age, members of this Mother-Reared (MR, N=6) monkey group were separated from their birth mothers and brought to the Yerkes Main Station in Atlanta, GA. There, they were progressively socialized in three peer groups of two home-cage partners and housed in adjoining 1.3m^3^ cages fitted with removable dividing panels.

Within forty-eight hours of birth, Nursery-Reared (NR, N=6) monkeys were randomly assigned to permanent separation from their mothers and hand-reared (i.e., diapered and bottle-fed) by animal care personnel, for approximately 90 days, at the YNPRC Main Station. Except for regular bottle-feeding, NR animals were continuously housed in adjoining 1.3m^3^ cages fitted with removable dividing panels, until 3 months of age. Then, at approximately 3 months of age, the NR group was separated into three peer groups of two home cage partners and housed in adjoining 1.3m^3^ cages. Thus, initial peer co-housing occurred at one year of age for MR (i.e., at maternal separation) and three months of age for NR (i.e., at cessation of bottle-feeding) subjects. All animals had olfactory, visual and auditory contact with conspecifics; but only co-housed peers had physical contact. Subjects received equal, ad libitum access to a standard diet of laboratory monkey chow as well as periodic fruits, vegetables, and nuts. All subjects were in a healthful condition when sacrificed for these experiments.

### Tissue Collection and Preparation

When the animals were approximately 1½ years of age, brain specimens were harvested, post-mortem subjects, for autoradiographic analysis. Following the intra-muscular administration of ketamine hydrochloride (10 mg/kg), each subject was sacrificed using an overdose of sodium pentobarbital, in keeping with the recommendations of the American Veterinary Medical Association Panel on Euthanasia. All post-mortem tissue assays were conducted in the laboratory of Dr. Paul Plotsky, at the Psychiatry Department of Emory University Medical School. Each brain was immediately extracted, washed in cold phosphate buffered saline (PBS) and cut into 1cm coronal blocks, using a rhesus brain matrix (A.S.I. Instruments, Warren, MA). The blocks were placed on an aluminum plate, over dry ice, and quickly frozen. They were subsequently stored at -80°C until sectioning on a cryostat.

On the day of cryosectioning, the tissue blocks were transferred, from -80°C storage, to dry ice. Using Tissue-Tek® Optimum Cutting Temperature Compound for cryostat sectioning (Sakura Finetek U.S.A., Inc., www.sakura.com) and ensuring that the midline was perpendicular to the platform, each block was temporarily attached to a tissue platform, on dry ice. On a cryostat cooled to -18°C, sections of 20 µm thickness were cut and thaw-mounted onto either a 3’’ X 31.5’’ pre-coated gelatin and poly-L-lysine glass slide (Brain Research Laboratories, Waban, MA) or a 3’’ X 31’’ Superfrost Plus glass slide (Fisher Scientific, www.fisherscientific.com). Sections were mounted on slides, dried at room temperature with a stream of cool air and, placed in air-tight containers with indicating silica gel packets (as a desiccant) and stored in a -80° C freezer, until autoradiographic analysis.

### Dopamine Receptor-1

DRD1 binding autoradiography assays were conducted according to the described procedures (Bergson et al., 1995). Brain sections containing the (left) prefrontal cortex, as well as the whole striatum and NAcc harvested from subjects were removed from -80° C storage and thawed. Following a 2-minute fixation in 0.1% paraformaldehyde (pH 7.4), to preserve tissue integrity, the sections were rinsed twice with an incubation buffer comprised of 50mM Tris-HCl, 0.1% BSA (pH 7.4) for 10 minutes each. The sections were then incubated for 90 minutes, at room temperature, with 1nM [125I]-(+)-SCH 23982 (Supplied by Robert Speth, Washington State University) in 50mM Tris-HCl (pH 7.4), 1mM MgCl2, 120mM NaCl, 5mM KCl, 2mM CaCl2, 0.1% BSA buffer and 1uM of the 5-HT2/5-HT1C receptor antagonist Ritanserin (Sigma-Aldrich Co., www.sigmaaldrich.com). Ritanserin was added to displace nonspecific binding of the radioligand from those 5-HT receptor subtypes. A set of consecutive sections was incubated in the same buffer, with radioligand (but without ritanserin) and 1uM of SCH23390 hydrochloride (Tocris Cookson Inc., www.tocris.com), which displaces [125I]-(+)-SCH 23982 binding from both DRD1 and 5HT receptors, serving as the nonspecific binding condition (NSB).

Following the incubation, sections were rinsed twice in cold 50mM Tris-HCl (pH 7.4), for ten minutes each. After being dipped in cold ddH2O, the slide-mounted sections were immediately dried, using a cool stream of air, for approximately 15 min. Then, both autoradiographic 125I microscale standards (Amersham Life Science, Inc., www.amersham.com) and the [125I]-(+)-SCH 23982 labeled sections were apposed to Kodak BiomaxMR X-ray film (Kodak, www.kodak.com) for 1-3 days.

### Dopamine Receptor-2

DRD2 binding autoradiography assays were conducted according to the methods that have been previously described (Bergson et al., 1995), with slight modification. Slides containing the left prefrontal cortex, as well as the whole striatum and NAcc were removed from -80° C storage and thawed, at 4°C. Approximately 2 hours later, they were dried for 10 minutes, under vacuum, at room temperature. Following a 2-minute fixation in 0.1% paraformaldehyde (pH 7.4), the sections were rinsed twice with an incubation buffer comprised of 50mM Tris-HCl, 0.1% BSA (pH 7.4), for 10 minutes each. The sections were then incubated for 45 minutes, at room temperature, with 1nM ^125^I-Epidepride (Supplied by Jouko Vepsäläinen, University of Kuopio, Finland), in 50mM Tris (pH 7.4), 0.1% BSA, 1mM MgCl2, 120mM NaCl, 5mM KCl, 2mM CaCl2, and 0.1% Ascorbic acid. 0.2µM of the α2-adrenergic receptor antagonist, Idazoxan (Sigma-Aldrich Co., www.sigmaaldrich.com) was added to displace nonspecific binding of the radioligand from α2-adrenergic receptor subtypes. A set of consecutive sections was incubated in the same buffer, with radioligand (but without idazoxan) and 10µM of the non-specific binding agent (+)-Butaclamol (Sigma-Aldrich Co., www.sigmaaldrich.com), to serve as nonspecific binding condition (NSB).

Following the incubation, sections were rinsed twice in cold 50mM Tris-HCl (pH 7.4), for ten minutes each. After being dipped in cold ddH2O, the slide-mounted sections were immediately dried, using a cool stream of air, for approximately 15 min. Then, both autoradiographic 125I microscale standards (Amersham Life Science, Inc., www.amersham.com) and the ^125^I-Epidepride labeled sections were apposed to Kodak BiomaxMR X-ray film (Kodak, www.kodak.com) for 1-3 days.

### Analysis of Receptor Autoradiographic Data

Autoradiograms from the DRD1 and DRD2 binding assays were digitized and then analyzed using the software program NIH Image (http://rsb.info.gov/nih-image; Bethesda, MD). Upon visual examination of these images, robust expression of DRD2 in the CLA necessitated the inclusion of this region as another potential stress-biomarker. Multiple, serial sections from the left PFC block and hypothalamic block were quantified for each research subject, by a treatment blind technician. For DRD1, optical density readings from the autoradiograms were converted to dpm/mg tissue equivalents using standard curves derived from measurement of the ^125^I-microscales standards. Contrastingly, for DRD2 optical density measurements were used, instead of dpm/mg tissue equivalents, because the ^125^I-microscales standards were below detection levels on film. For both DRD1 and DRD2, specific receptor binding was obtained by subtracting nonspecific binding from a corresponding area in the white matter, which represents brain areas where DADR binding should not exist.

All data analyses were conducted in SPSS® 16.0 for Microsoft Windows® (SPSS Inc., www.spss.com). Figures and tables were created using either Microsoft Excel (Microsoft Corporation, www.microsoft.com) or GraphPad Prism 10 for Microsoft Windows® (GraphPad Software, www.graphpad.com). The data analyzed included two independent variables (Mother- and Nursery-Rearing) and sixteen dependent variables for each (DRD1, DRD2) receptor binding experiment. The dependent variables included mean DRD1 or DRD2 density within the: prefrontal cortex (dorsolateral, medial, orbital, and total PFC), caudate (left, right, and total caudate), putamen (left, right, and total putamen), nucleus accumbens (left, right, and total NAcc), and claustrum (left, right, and total CLA).

DRD1 and DRD2 binding in the various prefrontal and striatal areas examined were compared using repeated measures analysis of variance (RMANOVA) with rearing (NR and MR) as a between factor and individual brain section as a within subject factor. In each case, Bonferroni corrections were employed, to account for the number of multiple comparisons. A 1-Tailed test was used in the two instances (i.e., prefrontal DRD1 and DRD2) where a specific directional prediction was made. A preliminary Shapiro-Wilks test of normality revealed that DRD2 binding in the right CLA of MR subjects violated normality assumptions (p=0.009), justifying the use of a non-parametric test. Therefore, the Mann-Whitney U Test, for comparing DRD2 in the CLA.

## RESULTS

### Basal Ganglia

No group differences were detected for the binding of DRD1 in the caudate, putamen, NAcc, or total striatum [Table 1]. Likewise, no group differences were observed for DRD2 binding in the caudate, putamen, NAcc, or total striatum [Table 2].

**Table 1.** [125I]-(+)-SCH 23982 Binding for Dopamine Receptor-1. No group differences were detected between nursery-reared (NR) and mother-reared (MR) animals, in the levels of DRD1 binding within the caudate, putamen, nucleus accumbens (NAcc), striatum, dorsolateral- prefrontal cortex (PFC), medial PFC or total PFC.

| Brain Region | Mother-Reared |  | Nursery-Reared |  | F | Degrees of Freedom | Significance (p Value) |
| --- | --- | --- | --- | --- | --- | --- | --- |
|  | Mean DRD1 (dpm/mg) | SEM | Mean DRD1 (dpm/mg) | SEM |  |  |  |
| Dorsolateral PFC | 5345.15 | 1002.31 | 5069.52 | 1143.82 | 0.03 | (1, 11) | $p > .05$ |
| Medial PFC | 4998.12 | 1143.22 | 4402.17 | 1319.21 | 0.12 | (1, 11) | $p > .05$ |
| Total PFC | 34972.36 | 3197.41 | 34119.43 | 4002.59 | 0.03 | (1, 11) | $p > .05$ |
| Total Caudate | 61683.20 | 1574.36 | 61107.82 | 806.50 | 0.12 | (1, 10) | $p > .05$ |
| Total Putamen | 60961.23 | 1650.18 | 61346.54 | 753.22 | 0.05 | (1, 10) | $p > .05$ |
| Total Nucleus Accumbens | 52296.71 | 3165.58 | 52596.84 | 872.52 | 0.01 | (1, 10) | $p > .05$ |
| Total Striatum | 172810.00 | 5454.39 | 175050.00 | 1990.42 | 0.17 | (1, 10) | $p > .05$ |

**Table 2.** ^125^I-Epidepride Binding for Dopamine Receptor-2. Nursery-reared (NR) and mother-reared (MR) animals did not differ in optical density (OD) of DRD2 binding in the caudate, putamen, nucleus accumbens (NAcc), dorsolateral prefrontal cortex (PFC), orbital PFC, or total PFC.

| Brain Region | Mother-Reared |  | Nursery-Reared |  | F | Degrees of Freedom | Significance (p Value) |
| --- | --- | --- | --- | --- | --- | --- | --- |
|  | Mean DRD2 (OD) | SEM | Mean DRD2 (OD) | SEM |  |  |  |
| Dorsolateral PFC | 0.17 | 0.01 | 0.15 | 0.02 | 0.94 | (1, 11) | $p > .05$ |
| Orbital PFC | 0.18 | 0.01 | 0.16 | 0.02 | 0.61 | (1, 11) | $p > .05$ |
| Total PFC | 0.63 | 0.03 | 0.55 | 0.05 | 1.69 | (1, 11) | $p > .05$ |
| Total Caudate | 3.53 | 0.39 | 3.29 | 0.34 | 0.22 | (1, 10) | $p > .05$ |
| Total Putamen | 3.05 | 0.23 | 3.06 | 0.36 | 0.00 | (1, 10) | $p > .05$ |
| Total Nucleus Accumbens | 2.03 | 0.20 | 2.00 | 0.19 | 0.01 | (1, 10) | $p > .05$ |
| Total Striatum | 8.74 | 0.73 | 8.46 | 0.84 | 0.06 | (1, 10) | $p > .05$ |

### Prefrontal Cortex

No group differences were detected for binding of either DRD1 [Table 1] or DRD2 [Table 2] in the dorsolateral PFC or total PFC. Similarly, no group differences were observed for DRD1 binding [Table 1] in the mPFC or DRD2 binding [Table 2] in the orbital PFC.

DRD1 (dpm/mg) binding within the oPFC was significantly greater in MR than in NR monkeys (1-Tail, RMANOVA, F (1, 11) = 4.138, p<.05, η2 = 0.293) [Figure 1]. In addition, DRD2 (OD) binding within the mPFC was significantly greater in MR, compared to NR monkeys (1-Tail, RMANOVA, F (1, 11) = 3.648, p<.05, η2 = 0.267) [Figure 2].

**Figure 1.**
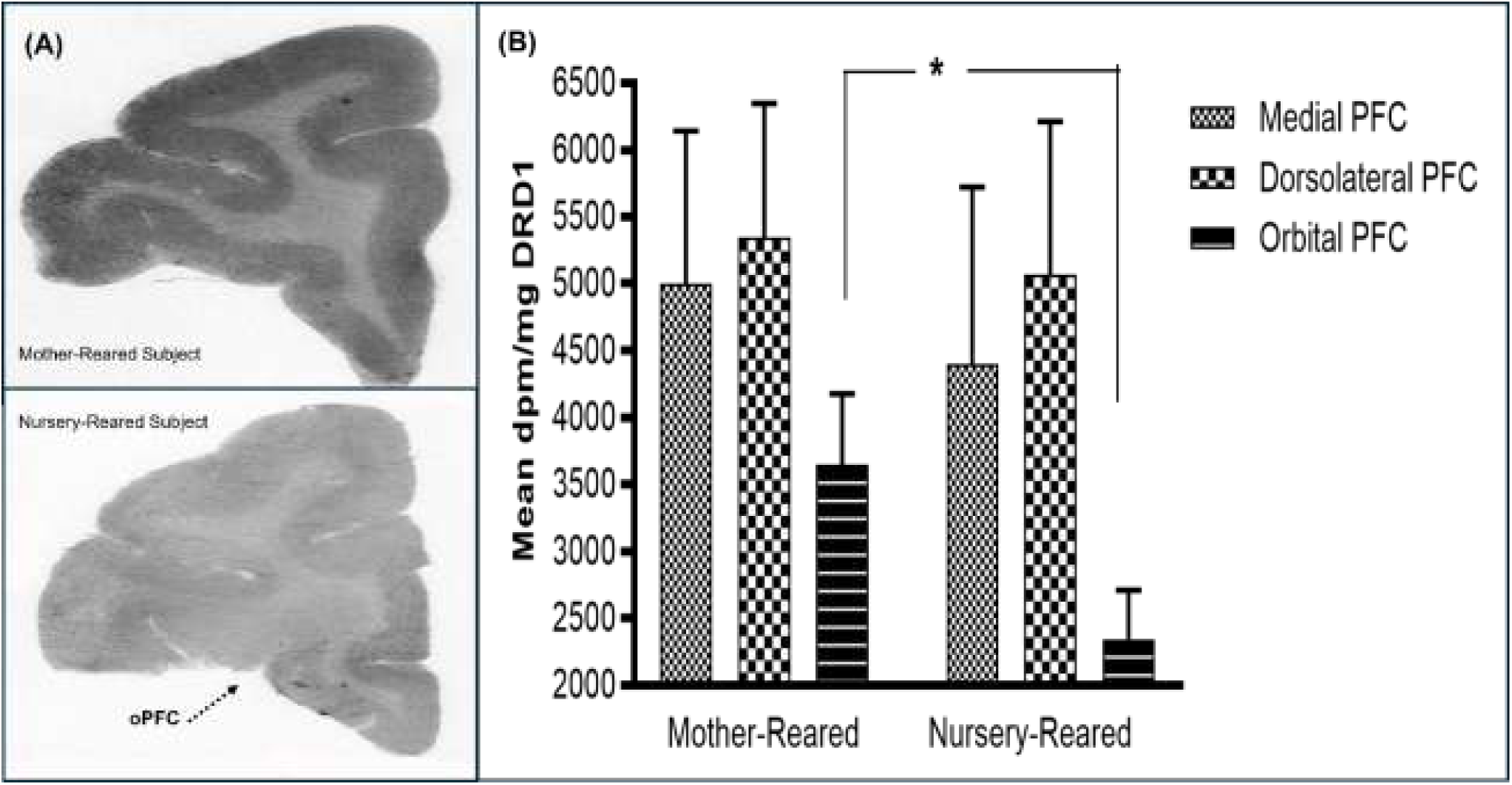
Reduced Dopamine Receptor-1 in the Orbital Prefrontal Cortex of Nursery-Reared Monkeys. *(A):* A pair of rhesus monkey autoradiograms from one mother-reared (MR) subject “RMC7” and one nursery-reared (NR) subject “RGZ6” illustrate [125I]-(+)-SCH 23982 binding for Dopamine Receptor-1 (DRD1) within the orbital prefrontal cortex (oPFC, indicated by arrow in bottom image). *(B)*: While not significantly different in the medial or dorsolateral areas of the prefrontal cortex monkeys significantly differed in the degree of DRD1 (dpm/mg) binding within the oPFC. MR (N=6) monkeys possessed greater DRD1 than NR (N=6) monkeys (1-Tail, RMANOVA, F (1, 11) = 4.138, p<.05, η2 = 0.293).

**Figure 2.**
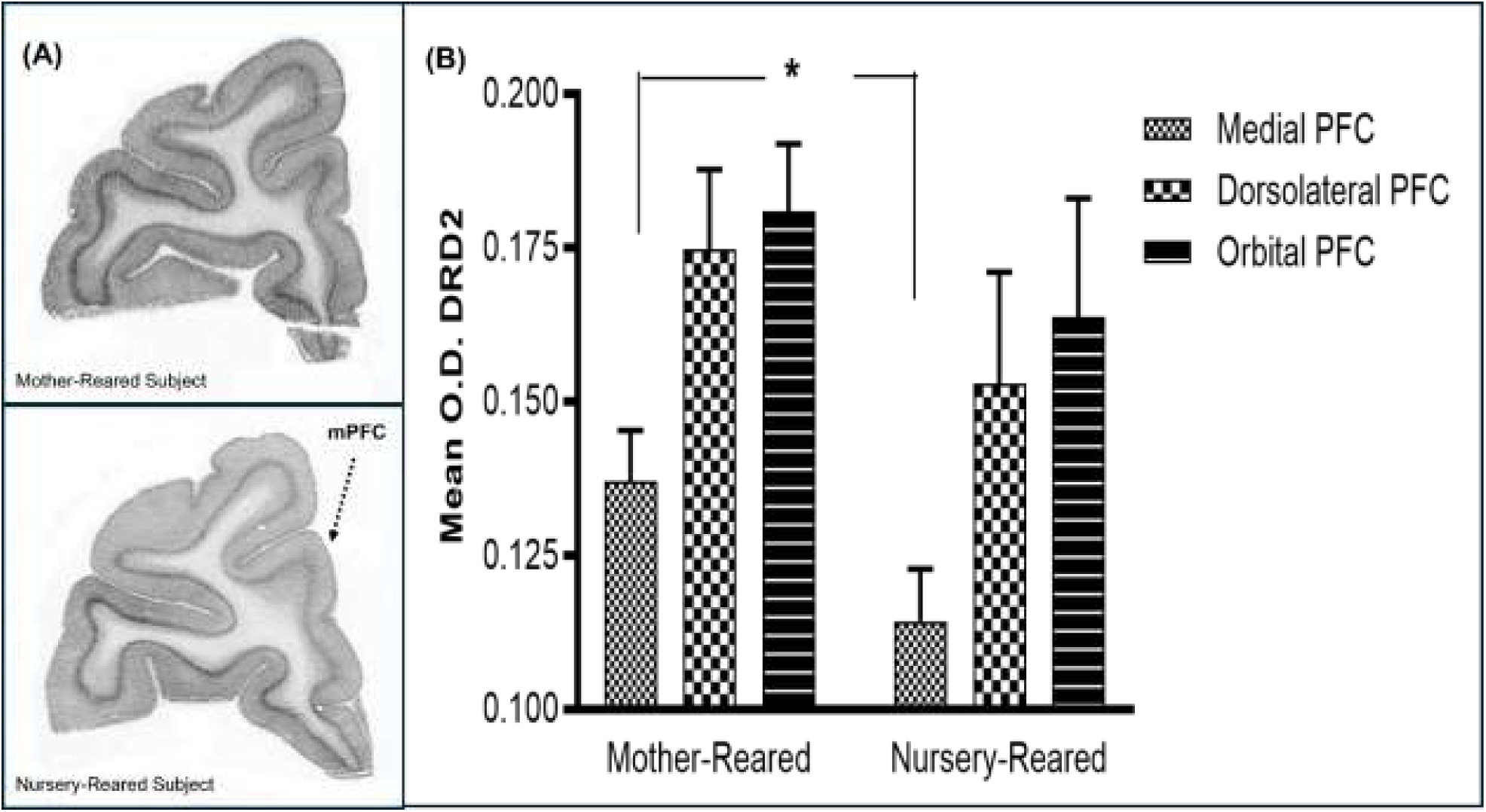
Reduced Dopamine Receptor-2 in the Medial Prefrontal Cortex of Nursery-Reared Monkeys. *(A)*: A pair of rhesus monkey autoradiograms illustrate ^125^I-Epidepride binding for Dopamine Receptor-2 (DRD2) in the medial prefrontal cortex (mPFC, indicated by arrow in bottom image) for one mother-reared (MR) subject “RMC7” and one nursery-reared (NR) subject “RRV6”. *(B)*: While not dissimilar in the dorsolateral or orbital prefrontal cortical regions, differently reared monkeys significantly differed in the degree of DRD2 binding within the mPFC. MR (N=6) monkeys possessed greater DRD2 optical density (OD) compared to NR (N=6) monkeys (1-Tail, RMANOVA, F (1, 11) = 3.648, p<.05, η2 = 0.267).

### Claustrum

DRD1 expression was barely visible in the CLA and therefore not quantified. The DRD2 signal was comparatively robust. While differences in Total and Left CLA DRD2 were not significant, MR subjects showed significantly greater Right CLA DRD2 than NR subjects (Mann–Whitney U = 27.5, p = 0.028; HL median difference = 0.045; rank-biserial r = 0.83) [Figure 3].

**Figure 3.**
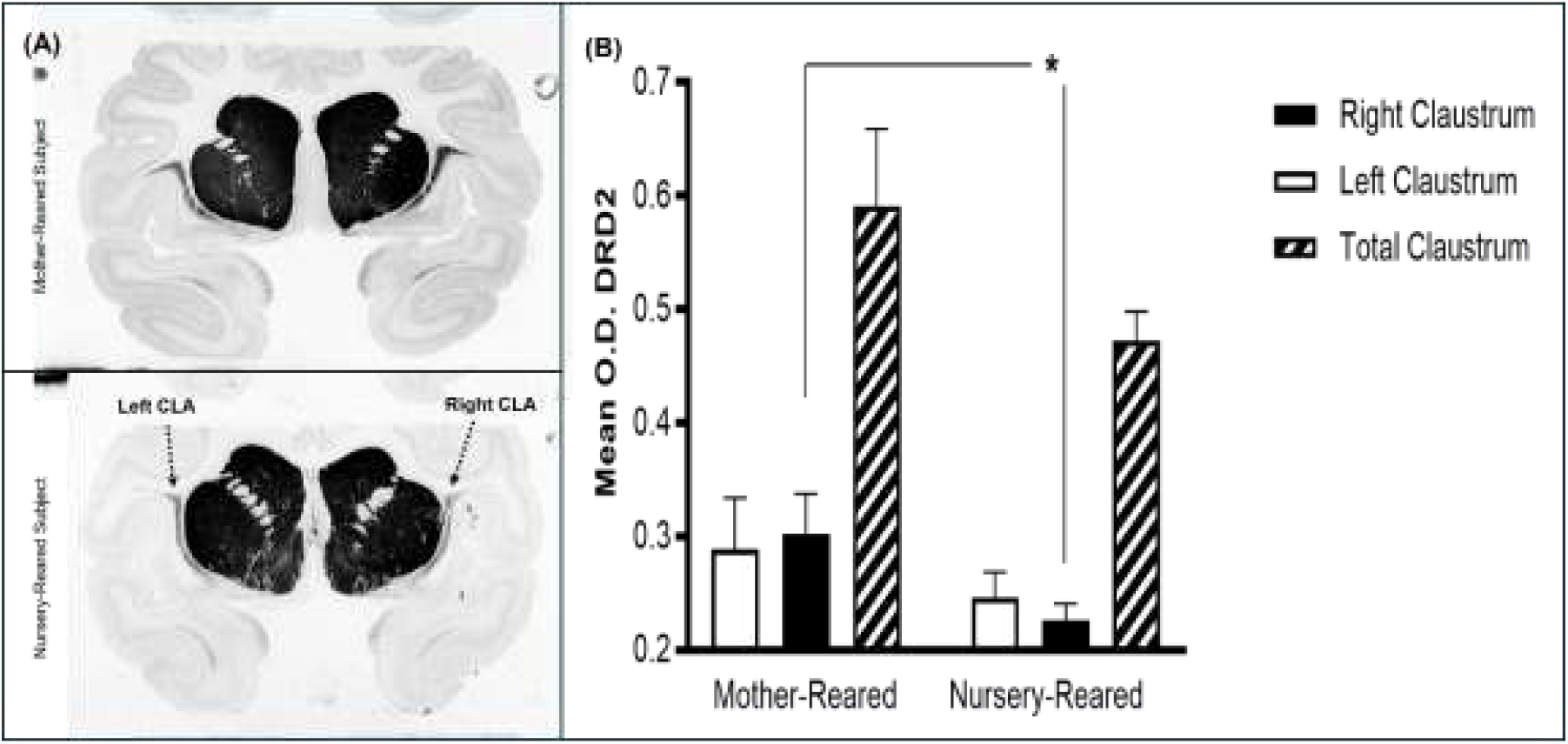
Reduced Dopamine Receptor-2 Binding in the Claustrum of Nursery-Reared Monkeys. *(A)*: Two rhesus monkey autoradiograms illustrate ^125^I-Epidepride binding for Dopamine Receptor-2 (DRD2) in the claustrum (CLA, indicated by arrows in bottom image) in one mother-reared (MR) subject “RMC7” and one nursery-reared (NR) subject “RGZ6”. (*B)*: Compared to NR (N=5) monkeys, MR (N=6) monkeys possessed significantly greater DRD2 optical density (OD) in the right claustrum (2-Tail, Mann–Whitney U = 27.5, p=<.05; HL median difference = 0.045; rank-biserial r = 0.83).

## DISCUSSION

In the current experiment, the impact of rearing on the neuroanatomical densities of DADR was examined through comparative receptor autoradiographic analyses on post-mortem brain specimens collected from 6 NR and 6 MR rhesus macaques. We first interrogated the possibility of *suppressed DA signaling* in NR by examining DRD1 and DRD2 densities in the basal ganglia. Our expected result of altered basal ganglia DRD1 and DRD2 in NR monkeys, compared to MR, was not realized. No group differences were detected for either the caudate, putamen, NAcc, or total striatum. Second, to address the possibility of weakened top-down cognitive suppression of limbic responses in NR, we *examined DRD1 and DRD2 densities in the PFC, with the prediction of decreased DRD1 and DRD2 in NR compared to MR monkeys.* Contrary to our expectations, there were no detectable group differences for either mPFC D1R, oPFC DRD2, or DRD1 and DRD2 in the dorsolateral PFC and total PFC. The hypothesized outcome whereby ELS of maternal deprivation and peer-rearing results in lower DRD1 and DRD2 in the PFC of NR compared with NR was realized. Our findings of 1) reduced oPFC DRD1 and 2) reduced mPFC DRD2 in NR compared with MR monkeys underscore the probability for suppressed DA signaling and weakened top-down cognitive suppression of limbic responses in NR. An unexpected finding of 3) decreased right CLA DRD2 binding in NR but not MR further corroborates a weakening of potential for top-down cognitive suppression of limbic responses in NR. In the text that follows, we will contextualize the negative results as well as the three noteworthy results and connected inferences in the broader literature on stress-related DA dynamics.

### I. Basal ganglia DRD1 and DRD2 may be impervious to maternal deprivation and peer-rearing

In their study, Martin et al. (1991) showed reductions in striatal TH immunoreactive axonal fibers, suggesting a reduction of dopaminergic neurotransmission in the striatum of isolation-reared monkeys. Instead, our data suggests region-specific effects of rearing for the PFC. Undoubtedly, the mPFC and orbital PFC (oPFC) differently modulate cortical DA output. While single-pulse action potential activation of the orbital PFC tends to inhibit firing in DA neurons, this increases DA activity when applied to the mPFC (Lodge, 2011). Consequently, nigrostriatal DRD1 and DRD2 may not completely mediate the increased propensities for abnormal and repetitive motor behaviors commonly associated with NR. Instead of representing simple pathologies, these tendencies may have self-soothing or vestibular stimulatory functions, as has been previously suggested (Kraemer et al., 1997).

### II. The PFC is particularly sensitive to maternal deprivation and peer-rearing

Together, the evolutionary newness of the PFC (Preuss, 1995), it’s protracted development trajectory (Gogtay et al., 2004), and connections with the HPA (Arnsten, 2015, 2009) dictate it would be more open to early insult than the basal ganglia. Over the course of their evolution, not only did the frontal lobe significantly expand (Gibson et al., 2001; Gould, 2001; Kaskan and Finlay, 2001; Rakik and Kornack, 2001), but the PFC developed structural heterogeneity in keeping with demands for behavioral and cognitive complexity among human and non-human primates (Cruz-Rizzolo et al., 2011). Simultaneously, the PFC’s immense complexity requires a balance of adequate benefits and minimal costs that outweigh the energy demands associated with its operation. It is not surprising, therefore, that extreme stress would result in compensatory declines in dopaminergic neurotransmission.

### III. Maternal deprivation and peer-rearing lower DRD1 in the orbital PFC

DA release in the PFC is intended to focus attention on reward-indicating stimuli or to predict reward itself (Heinz, 2002). In this regard, the oPFC is critically important for the estimation of reward value (Wallis and Miller, 2003). Although DRD1 and DRD2 cooperatively facilitate the flexibility of behavior, PFC DRD1 activity is primarily important for the execution of working memory functions (Floresco and Magyar, 2006). Here, DRD1 (dpm/mg) binding in the oPFC was found to be greater in MR than in NR monkeys. A loss of orbitofrontal PFC DRD1 indicates that changes in the dopaminergic modulation of HPA function may characterize known differences in emotional and motivational regulation between NR and MR monkeys. Diminished reward learning and impaired decision making are associated with ELS (Hanson et al., 2017; Sheridan et al., 2018; Weller and Fisher, 2013).

### IV. Maternal deprivation and peer-rearing lower DRD2 in the medial PFC

The finding of differences within the mPFC is consistent with known differences in stress physiology, behavior and emotion regulation between NR and MR monkeys. Specifically, they suggest that the altered attention and motivation (Sanchez et al., 2001) or inclination towards risky behavior (Sanchez et al., 2001) often observed in NR, compared to MR monkeys, can be attributed to altered mesocortical-mesoprefrontal DA neurotransmission. While mPFC DRD1 blockade (with SCH23390) reduces preference for large/risky options, mPFC DRD2 stimulation (using quinpirole) completely impairs decision making (St Onge et al., 2011). In our study, DRD2 binding was elevated in the mPFC of MR compared to NR and appeared most marked in area 32 of the mPFC. This is intriguing because, as part of the prelimbic cortex, area 32 has connections with the amygdala (Majak et al., 2002) and participates in autonomic control (Cechetto and Chen, 1990; Oppenheimer and Cechetto, 1990) through synapses in the lateral hypothalamus. The binding also appeared more prevalent in layer 5, which contains pyramidal cells and is thus involved in motor output. The PFC, and particularly the medial portion, is highly sensitive to stress (Crane et al., 2003; Figueiredo et al., 2003; Finlay et al., 1995; Jedema et al., 1999; Stevenson and Gratton, 2003). For example, acute stress has been shown to trigger release of NE and DA within the mPFC (Finlay et al., 1995; Jedema et al., 1999), which can in turn suppress the HPA response to stress (Crane et al., 2003).

Also, in rats, lesions within the mPFC can result in increased adrenocorticotrophin and corticosterone production following restraint stress (Diorio et al., 1993). As a biomarker for stress related depression, childhood maltreatment is linked to greater connectivity between the ventral striatum and the mPFC (Hanson et al., 2018). Theoretically, initially enhanced DA neurotransmission could be followed by compensatory reductions in DRD2.

### V. Maternal deprivation and peer-rearing decrease right claustrum dopamine receptor-2

The CLA has alternatively been described as mysterious (Goll et al., 2015), enigmatic (Crick and Koch, 2005), and the “ “gate keeper” of neural information for consciousness awareness” (Torgerson et al., 2015). Although its properties are yet to be fully described, reciprocal connections with midbrain dopaminergic neurons, the basal forebrain, and amygdala strongly suggest a role in attention regulation, inhibition, sensory-emotional integration, and susceptibility to cocaine (Atlan et al., 2018; Chen et al., 2023; Jackson et al., 2020; Reser et al., 2017; Zhao et al., 2024). Notably, in a study of Arc-dVenus reporter mice, a circuit linking the CLA with neurons of the basolateral amygdala can produce anxiety-like behaviors when optogenetically photostimulated, and increase resilience to chronic social defeat stress when deactivated (Niu et al., 2022).

With attention deficits, impulsivity, addiction vulnerability, and anxious tendencies, another common product of ELS is increased susceptibility to chronic pain (Krantz et al., 2019). In male and female C57BL6/J mice, pain learning is mediated by neural pathways bridging the anterior cingulate cortex and claustrum, which can be impaired by chronic inflammatory pain (Faig et al., 2024). In a recent fMRI study of humans, cognitive difficulties associated with chronic pain were mediated by connections between the CLA and right dorsolateral PFC (Stewart et al., 2024).

Although this study only examined the left PFC, it is conceivable that an ipsilateral loss of DRD2 in the right dorsolateral PFC or anterior cingulate could be observed in a future experiment. Finally, evidence that the CLA plays an important role in inhibiting the PFC (Jackson et al., 2018) underscores our analysis that the loss of DRD2 in NR functionally weakens top-down cognitive suppression of limbic responses. A Hodges-Lehmann (HL) median difference, which is a robust estimate of group divergence, indicated that MR subjects typically score about 0.045 higher than Nursery on Right DRD2.

The psychobiological sequelae of early trauma are determined through complex interactions between social environmental and genetic factors shaping behavior as well as the cytoarchitecture and molecular landscape of the brain. Epigenetic modifications to DA receptor genes may precipitate behavioral and pharmacological sensitivities in response to early stress. For example, in a study of DRD2 CpG methylation and personality traits among 51 differently reared chimpanzees, Staes and colleagues (2022) observed extroversion to be significantly linked to methylation in NR but not MR subjects. In keeping with an “experience-dependent methylome”, this suggests DA receptor genes are sensitive to maternal deprivation. Conversely, in a study of 279 Dutch adolescents, a predisposition to develop emotional eating subsequent to early adverse rearing was linked to a possession of an A1 allele for the DRD2 gene Taq1A polymorphism (rs1800497), which is itself associated with attenuated brain DRD2 receptor availability (van Strien et al., 2010). It is not known on what genetic background unfolded the interplay between rearing and DADR in our experiment. Nevertheless, our controlled experiment on randomly assigned research subjects produced findings that cannot be explained by either the diathesis-stress or differential susceptibility hypotheses. While genetically determined pre-existing vulnerabilities or increased tolerance to stress can play a role, as in multigenerational studies, the current findings are best explained using an evolutionary developmental and ecological framework that accommodates experience. Although this is a likely explanation, it is unknown whether the reductions in DADR densities observed in NR monkeys represent a downregulation of receptors, in consequence of increased dopaminergic transmission in the PFC. Alternatively, the receptor downregulation could be a result of group differences in DADR gene expression or divergent transcriptomes. Both possibilities represent new avenues for future research.

Despite the relevance of dopaminergic circuits for understanding the impact of early life stress, there has been insufficient progress towards characterizing the neurophysiological alterations and determining future risk for mental health disorders (Smith and Pollak, 2020). In a prior publication, we showed that the impact of ELS in the form of maternal deprivation is lower DA tone that is modulated altered HPA function (Seraphin et al., 2022). The current study demonstrates that dopaminergic circuits respond to early maternal deprivation and peer rearing by compensatory reductions of DRD1 in the orbital PFC, and DRD2 in both the mPFC and CLA. These results suggest a “hypodopaminergic prefrontal-claustral ecophenotype model” [Figure 4] wherein the early stress of maternal deprivation and peer rearing cause compensatory neuroendocrine responses that decrease dopaminergic function and eventually culminate in a behavioral profile commonly associated with early maltreatment. While the net effect is a diminished ability to inhibit impulses and increased limbic irritability, these outcomes should be considered adaptations to extraordinarily challenging developmental conditions.

**Figure 4.**
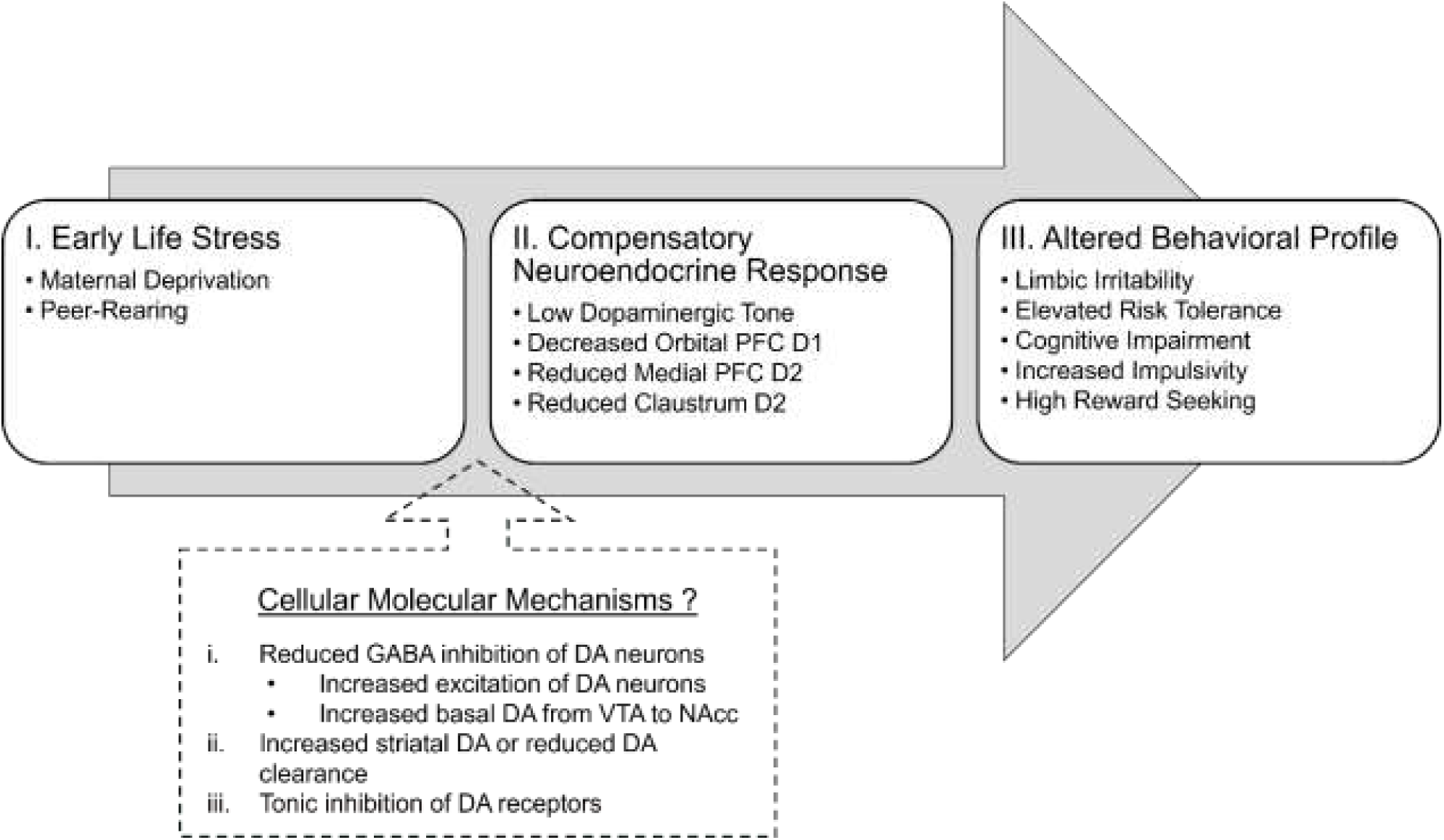
Hypodopaminergic Prefrontal–Claustral Ecophenotype Model. According to our hypodopaminergic prefrontal-claustral ecophenotype model, early stress leads to compensatory neuroendocrine responses that reduce dopaminergic function and culminate in altered behavior.

## ACKNOWLEDGEMENTS

This investigation was part of the dissertation research of S.B. Seraphin in the Department of Anthropology and Center for Behavioral Neuroscience at Emory University in Atlanta, Georgia. This work is being published after the passing of major contributors, Dr. James Timothy Winslow (J.T.W.) and Dr. Patricia L. Whitten (P.L.W). The research was supported by NIH grants MH57704 (J.T.W) and NIH base grant RR-00165 to the Yerkes National Primate Research Center (YNPRC), as well as by a National Alliance for Research on Schizophrenia and Depression (NARSAD) Young Investigator Award (M.M.S.), and a Center for Behavioral Neuroscience venture grant (STC Program, NSF, under agreement No. IBN-9876754; J.T.W., M.M.S., P.L.W.). The YNPRC is fully accredited by the American Association for Accreditation of Laboratory Animal Care. The authors are grateful for the generous support of Robert Speth Ph.D., Jouko Vepsäläinen, Ph.D., Paul Plotsky, Ph.D., Todd Preuss, Ph.D., Pamela Noble, Casie Lyon, Thomas Insell, M.D., and Larry J. Young, Ph.D.

